# From seven combination hypotheses to one testable interaction: a gated agentic AI–QSP workflow applied to healthy-ageing interventions

**DOI:** 10.64898/2026.09.29.755460

**Authors:** Igor Goryanin, Bob Damms, Irina V. Goryanin

## Abstract

Large language model (LLM) agents can propose combination therapies and construct supporting mechanistic models far quicker than either can be verified. To address this gap, we built a gated agentic AI–quantitative systems pharmacology (QSP) workflow where no hypothesis reaches a report until it clears strict hurdles for precedent, evidence, structural integrity, and release. We applied this pipeline end to end to healthy-ageing interventions. It began with a 34-state model calibrated on clinical trial data for semaglutide and metformin. Locked and evaluated against an independent post hoc DNA-methylation trial, the model captured the direction of change across all 9 organ and system clocks, but the magnitude was far too small: it predicted about −0.14 years per year where −1.6 to −5.0 were observed, halting tier promotion. The workflow then generated seven nutritional combination hypotheses (H1–H7) with 48 new parameters encoded in SBML. Identifiability analysis showed that 37 of these were completely unconstrained by available data. Trimming the model down to its lead pairing—a food-derived bioactive (Agent A) plus a single-strain probiotic (Agent B)—left six identifiable parameters and locked the uncalibrated interaction term at zero. From this reduced model, we derived an 84-day $2 \times 2$ factorial trial with a 28-day washout, requiring 356 participants for 80% power at an interaction ratio of 0.75. A 9,000-person virtual trial confirmed the design (80.5% power), but revealed an operational trap: an assay floor at 24 mg/kg slashed power to 33% and skewed estimates toward the null, forcing a protocol-level censoring condition before study lock. Testing the pipeline across downstream operational stages (10–42) and comparative oncology programmes showed that unconstrained generation consistently over-reports novelty, topological entry points can masquerade as biological mechanism, and governance gates remain vulnerable to operator overrides. Ultimately, the workflow delivered no silver bullet: its output is a single testable interaction, an assay-hardened trial design, and an audit trail documenting why the other six hypotheses were shelved. (Agent identities, doses, and agent-specific sources are withheld in this version pending patent filing).

## Introduction

Healthy-ageing interventions are increasingly tested in combination. Single agents produce small effects on ageing biomarkers, and additive signals have been reported when interventions are combined, as in one large factorial trial of supplements and exercise (reference withheld). The number of candidate combinations, doses and schedules grows far faster than any trial programme can explore. Mechanistic quantitative systems pharmacology (QSP) models have long been proposed for ranking such options before they reach the clinic [1,2].

Two developments change how that ranking is done. Large language models (LLMs) can now read a literature corpus, propose targets and pairings, and draft model code to test them within hours [3]. A recent benchmark of frontier models on ageing-biology tasks found that no single model dominated and that performance shifted with how questions were phrased [4]. Generation is no longer the scarce step; checking is.

An AI that proposes combinations fails in characteristic ways. It over-reports novelty, because it reasons from the corpus it was given rather than the world. It builds models that reproduce the target phenomenon for reasons unrelated to the stated mechanism, and adds parameters that no available data can identify. It ranks options by artefacts of model structure, and writes fluent claims that outlive the evidence behind them. None of these failures is visible in the output; all of them are visible in an audit trail, if one is kept.

We previously used a QSP model of metabolic and ageing pharmacology to compare semaglutide- and metformin-based combinations [5]. Here we ask a different question: what happens when an agentic workflow generates new combination hypotheses, builds and calibrates the models, and designs the trial, under explicit gates? We report (i) the workflow at architecture level; (ii) the complete record of one ageing programme, AgeQSP, from its semaglutide and metformin branches through seven hypotheses built on two coded nutritional agents to a single testable interaction; (iii) a model-derived factorial design and virtual trial; (iv) the point at which the workflow stopped, and why; and (v) how the same gates behaved in two other programmes. To protect pending intellectual property, the identities, doses and agent-specific sources of the combination partners are withheld in this version and replaced by codes (Methods); the workflow, model structure, identifiability analysis, trial design (without doses) and virtual-trial results are reported.

## Results

### A gated workflow separates generation from reporting

In the workflow, anything may be generated, but a hypothesis reaches a report only through three gates (Figure 1). The AgeQSP programme ran under version 2.21 of the workflow specification, which divides the work into numbered stages from intent (0X) through literature intelligence (0.0), model construction and export (0A–0L), hypotheses (3), calibration (5), validation (6), sensitivity and identifiability (7–8), evidence synthesis (9), virtual populations (9A), combination analysis (10) and publication control (11–13).

**Figure 1.**
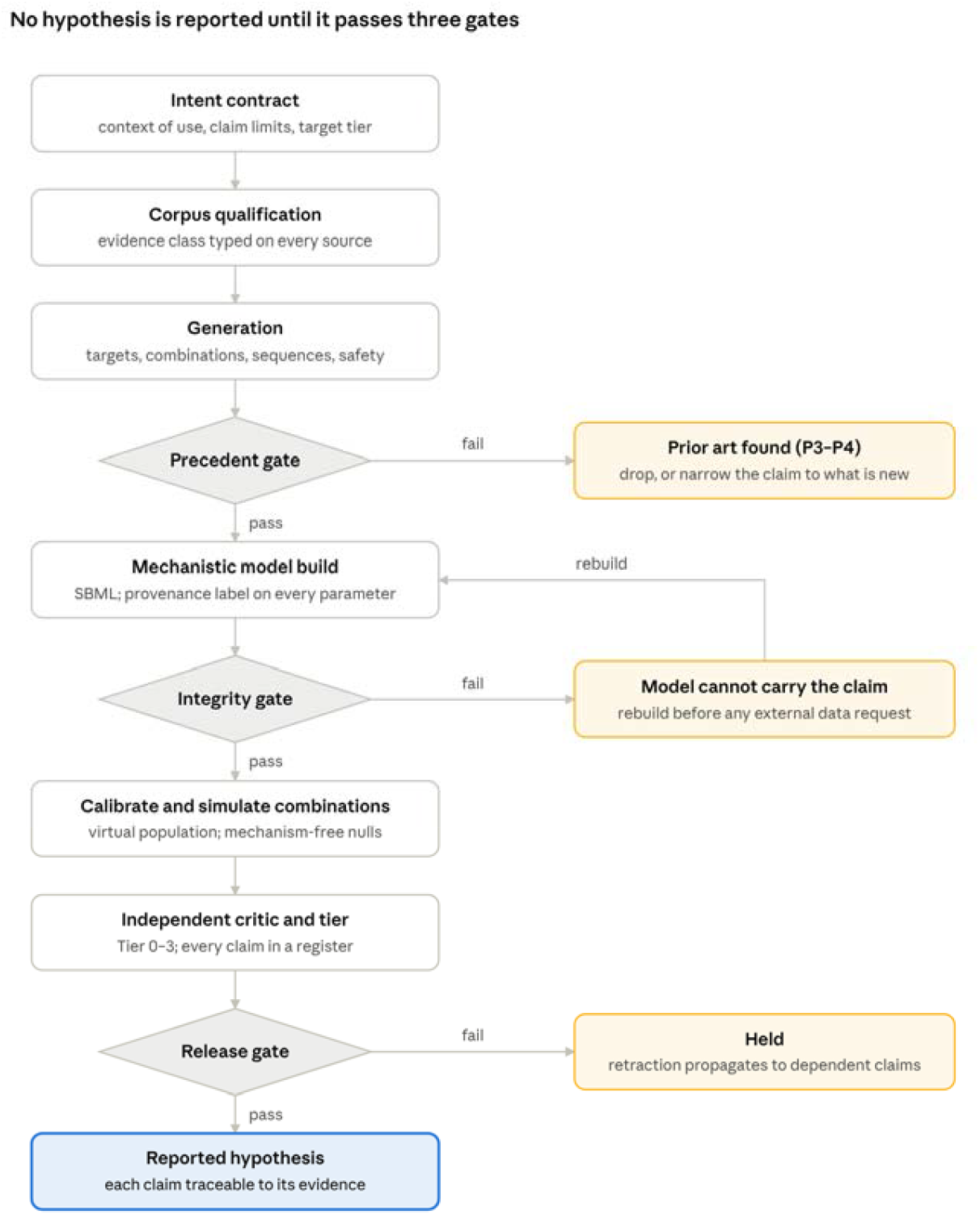
The gated workflow. Hypotheses may be generated freely but reach a report only through a precedent gate, a model-integrity gate and a release gate. Every failure exit is logged, so the effect of each gate on the final list can be counted. Box 1 describes how the gates were implemented in AgeQSP.

Every failure exit is logged, so the effect of each gate on the final list can be counted rather than asserted.

Each stage must end with exactly one verdict line: PASS, HOLD or BLOCKED. PASS is allowed only when the stage’s work and gate checks are complete, and the specification forbids advancing past a failed gate. The operator advanced the pipeline with short commands (“next stage”, “continue”, “perform all next stages”), so the stopping points reported below were set by the gates, not by the operator.

Three further controls run throughout. Every model file declares a state (for example EXTENSION_ENCODED or EXTENSION_PARTIALLY_CALIBRATED_MONOTHERAPY), and some transitions are forbidden, such as moving an extension straight to a calibrated core. Every parameter carries a provenance label, such as ASSUMED_SENSITIVITY_ONLY, FITTED_SURROGATE or FITTED_FIGURE_DIGITIZED_AGGREGATE. And a 22-item critic checklist (A1–A22) is re-run after every change to the model file. Models are encoded in SBML Level 3 [6] and packaged with SED-ML in COMBINE archives [7], and every result carries a validation tier in the spirit of risk-informed credibility frameworks [8,9].

#### Box 1. How the three gates in Figure 1 were implemented in AgeQSP

| Gate | Implementing stages | What it checked in AgeQSP |
| --- | --- | --- |
| Precedent and evidence | 0.0 literature intelligence; evidence ledger; contradiction–support map | Direct combination evidence for each hypothesis; conflicts between sources; whether any data could calibrate an interaction |
| Model integrity | 0B-A compartment verification; 0D-A rate-law equivalence; 0H dimensional audit; 0J and 0K replay and export; critic A1–A22; 7–8 sensitivity and identifiability | Structure, units and executable equivalence; whether parameters are identifiable from feasible observations |
| Release | 11–13 publication control; 12A claim-warning table; model locks; hash-chained stage manifests | Prohibited wording, validation tier, and whether any synthetic result could be presented as evidence |

The precedent gate in AgeQSP was lighter than in later versions of the specification: it recorded direct combination evidence per hypothesis but did not assign formal precedent classes or search patents systematically.

### AgeQSP: drug branches calibrated, clock predictions failed

AgeQSP went through twelve steps with a recorded verdict at each (Figure 2). Its first version answered a structural question: could a latent-state model separate ageing biology from the measurement processes of individual clocks, with outputs limited to research and trial design? It had 31 states on a 0–1 scale: nine systemic ageing processes (inflammaging, senescent burden, mitochondrial dysfunction, proteostasis, nutrient sensing, vascular dysfunction, immune decline, stem-cell decline and frailty) and damage and reserve for each of 11 organs. Of its 60 parameters, 56 were assumed. Calibration was blocked because no longitudinal subject-level data were available.

**Figure 2.**
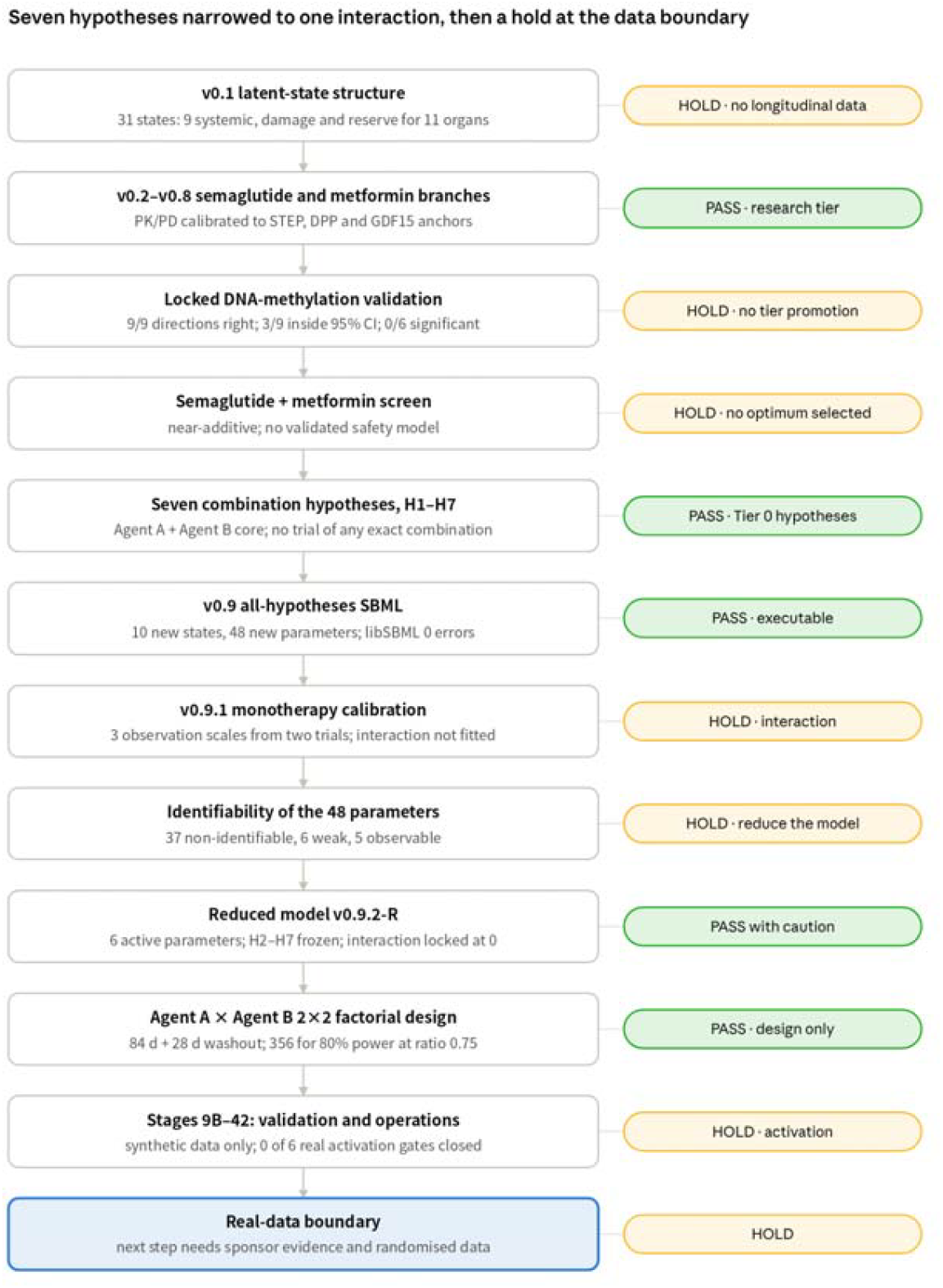
Development path of AgeQSP. Twelve steps from the first latent-state structure to the hold at the real-data boundary, with the verdict recorded at each step (green, PASS; amber, HOLD). Seven hypotheses built on Agents A and B were narrowed to one interaction, whose coefficient stays locked at zero until randomised factorial data exist.

Versions 0.2–0.8 added drug branches. A surrogate observation layer fitted six parameters to nine organ-age contrasts from 185 CALERIE participants (in-sample RMSE 0.038 years); the mechanistic core was not fitted [10]. Versions 0.3 and 0.4 added subcutaneous semaglutide (two-compartment pharmacokinetics, GLP-1 receptor engagement) and oral metformin, with GDF15 and AMPK mediators, for 34 states. Six semaglutide parameters were calibrated to published STEP 1, 4 and 5 weight trajectories, pooled CRP ratios and HOMA-IR [11–14], giving a weight RMSE of 1.60 percentage points and a modelled CRP ratio of 0.562.

The locked semaglutide model was then tested against an independent post hoc DNA-methylation analysis of a 32-week randomised trial in 84 participants with HIV-associated lipohypertrophy [15], with nothing refitted (Figure 3). The model’s HIV and antiretroviral population offsets were set to zero, so transportability is itself a limitation of this test. All nine organ and system endpoints moved in the observed direction. Only three (lung, immune and musculoskeletal) fell inside their 95% confidence intervals, and none of the six statistically significant effects did: the model predicted about −0.14 years per year where −1.60 to −4.99 were observed (RMSE 3.68). Exploratory clock mappings failed similarly; PhenoAge, for example, was predicted at −0.22 against −4.9. The stage returned HOLD and the validation tier was not promoted.

**Figure 3.**
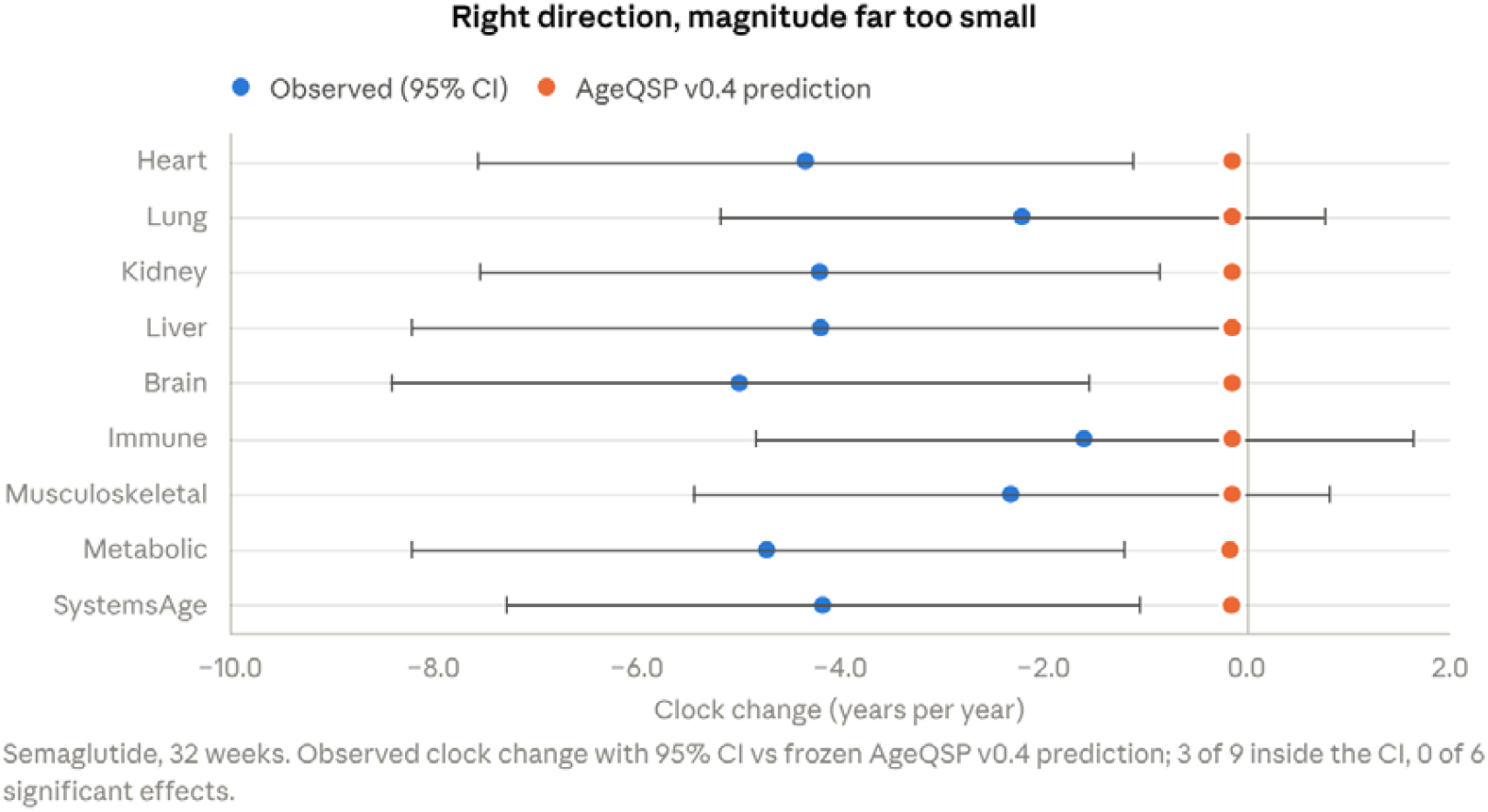
Locked external validation of the semaglutide branch against a post hoc DNA-methylation analysis of a 32-week trial in 84 participants with HIV-associated lipohypertrophy [15]. Blue, observed annualised clock change with 95% CI; orange, frozen AgeQSP v0.4 prediction. Direction agreed for all 9 endpoints; 3 of 9 predictions fell inside the CI and none of the 6 significant effects did. Source: Stage 6R.

The failure is structural, not parametric. Global sensitivity analysis (3,000 Latin-hypercube samples, ±30% on 20 parameters) placed the 5th–95th percentile of the whole-system effect at −0.208 to −0.094 years per year, far from the observed −4.17. Its top driver, the inflammation clearance rate (Spearman ρ −0.615), is itself an assumed core parameter.

The other analysis stages behaved as specified but added little evidence. The two inflammation terms were perfectly correlated (r > 0.9999999999), so the fit was rank 5 of 6 until they were merged; after repair, four active parameters had relative standard errors of 11–31%. A Bayesian synthesis of STEP 1–3 gave a weight effect of −9.61 percentage points (95% CrI −15.19 to −3.98; I^2^ 97.3%) and a CRP ratio of 0.564 (0.472–0.674), which is not independent of the calibration. A 9,000-person virtual population passed 39 of 39 aggregate checks but carries no individual-level claim.

A screen of semaglutide plus metformin behaved almost additively: interaction residuals were of order 10^−5^ on the model’s inflammation, nutrient-sensing and system-age states. After the metformin GDF15 driver was moved from plasma to gut exposure, to match human data [16], the combination added weight loss (−16.00% vs −13.98% for semaglutide alone) and a small HOMA-IR proxy gain (0.673 vs 0.680) but left the CRP proxy unchanged (0.565). The GDF15 fold at 12 weeks remained discordant with the AIM study (model 1.86, observed 1.35) [17]. With no quantitative safety model and no validated ageing endpoint, the stage returned HOLD and selected no optimum.

The publication gate then removed 15 prohibited phrasings from the draft report, including “accurately predicts biological age”, “optimal anti-aging combination” and “SBML/COMBINE-compliant”, the last because formal SBML validation had not been run.

### Seven combination hypotheses, and the evidence behind each

The operator then directed the workflow towards gut–immune combinations built on a food-derived bioactive ingredient (Agent A) and a single-strain probiotic (Agent B). The literature stage generated seven hypotheses (Table 1), each with a mechanism, a primary prediction, a falsifier and a statement of direct combination evidence, and scored each from 1 to 5 on evidence strength, mechanistic orthogonality, safety and feasibility, and development priority. No trial of any exact combination was found.

**Table 1.** All combination hypotheses generated (full text and sources in Table S1) E, evidence strength; O, mechanistic orthogonality; S, safety and feasibility; P, development priority. Frozen: excluded from active fitting in AgeQSP v0.9.2-R.

| ID | Combination | Predicted effect | Falsified if | Direct combination evidence | E/O/S/P | Status |
| --- | --- | --- | --- | --- | --- | --- |
| H1 | Agent A + Agent B, at the daily doses of their monotherapy trials | Faecal calprotectin and hsCRP/IL-6 fall more than with either alone; T-cell response kept | No AxB interaction and no microbiome/SCFA mediation, or worse tolerability | None in older adults; indirect evidence from one small crossover study of a related pairing (reference withheld) | 4/4/5/5 | Lead: reduced model and trial design |
| H2 | H1 + Agent C (a widely used nutritional supplement) | Larger fall in inflammatory burden and DNA-methylation clock measures than Agent C alone | No gain over Agent C on prespecified endpoints | None | 4/5/4/5 | Frozen; add-on only if the H1 interaction replicates |
| H3 | H1 + Agent D (a supplement with proposed mitochondrial effects) | Muscle endurance, mitochondrial markers and gut inflammation improve beyond either axis | No cross-axis gain, or no functional translation | None | 3/5/4/4 | Frozen; add-on only if the H1 interaction replicates |
| H4 | An incretin-class drug + H1, older adults with obesity | Less residual inflammation at matched weight loss | No residual benefit after weight adjustment, or excess GI intolerance or infection | None | 3/4/3/4 | Frozen; exploratory |
| H5 | A biguanide + H1 | Gut-inflammatory change beyond the drug's metabolic effect; no weight synergy assumed | No separable gut benefit, or additive GI intolerance | None; mechanisms overlap | 2/3/3/3 | Frozen |
| H6 | Agent E (an immunomodulatory drug class; drug and dose not fixed) + H1 | Better antiviral immune response | No clinical infection benefit despite biomarker gains | None | 2/4/2/2 | Frozen; lower priority |
| H7 | Agent A + Agent F (a synbiotic) + Agent G (a micronutrient) | More target-taxon abundance and SCFA capacity; efficacy not inferred from taxa alone | Taxa shift without metabolite, barrier or inflammatory gain | A small related multi-component study shifted taxa without metabolite, barrier or inflammatory gain (reference withheld) | 3/3/5/3 | Frozen; doses not fixed |

The evidence behind the hypotheses was thin and, on the systemic endpoints, contradictory. In a short randomised trial, Agent A improved ex vivo immune-cell readouts, but systemic IL-6 and CRP were not lower than with placebo (reference withheld). In a short randomised trial, Agent B lowered faecal calprotectin, with a non-significant CRP trend (reference withheld). A meta-analysis of 29 probiotic, prebiotic and synbiotic trials in 1,633 older adults was too heterogeneous to serve as a calibration target [18]. Eight entries in a contradiction–support map (Table S2) recorded these conflicts, and the last of them, the absence of any factorial A×B data, carried the verdict BLOCK CALIBRATION.

The evidence gate changed the design before any model was fitted. The first factorial draft had faecal calprotectin and hsCRP as co-primary endpoints. Because neither monotherapy moved systemic CRP, the ledger’s instruction was not to force a systemic effect, and hsCRP was moved to an exploratory endpoint. The stage also set an advancement rule: an Agent C or Agent D add-on (H2, H3) is tested only if H1 first shows a reproducible interaction.

### Encoding all seven hypotheses exposed what the data could not identify

All seven hypotheses were encoded in one SBML extension, AgeQSP v0.9, as switchable branches on the validated parent. It added 10 states (Agent A gut exposure, Agent B and Agent F engraftment, a prebiotic substrate, SCFA, gut repair, and Agent C, D, E and G signals), 8 switches, 4 exploratory observables and two gut compartments. Six parent rate rules gained multiplicative terms; 62 were unchanged. The interaction entered only through the engraftment term, dB/dt = k_on·(1 − B)·(1 + η·A_gut) − k_off·B, with η = 0.

The extension passed the executable gate. libSBML 5.21.2 reported 0 errors and 0 fatal findings (1,067 warnings, all unit-declaration code 99505), libSEDML 2.0.34 reported none across 7 scenario models, and all 84 cross-document targets resolved. With every new switch off, the extension reproduced the parent to a maximum relative difference of 1.5×10^−16^. Six error-level findings on the first pass came from duplicate annotation namespaces and were fixed without changing any equation.

Monotherapy calibration (v0.9.1) fitted three observation scales to three digitised targets from the two monotherapy trials: Agent A’s effects on a stimulated cytokine response and on T-cell counts, and Agent B’s effect on faecal calprotectin (target values withheld with their sources). With three parameters and three targets the fit is exact by construction; relative standard errors of 18–46% rest on assumed endpoint standard deviations. Two unfitted checks agreed with the trials: the model’s systemic inflammation ratio stayed at 0.999 for Agent B and 0.997 for Agent A. The critic held item A15, the interaction, which no monotherapy data can inform.

Identifiability then failed at scale. Across 48 extension parameters and 147 design observations the sensitivity matrix had full rank but a condition number of about 1.2×10^10^. Only 5 parameters, all observation scales, were locally observable (RSE 2.7–6.9%). Six, the Agent C, D and E on/off pairs, were weak or confounded (pairwise correlation above 0.95). The other 37 were non-identifiable, including every Agent A, Agent B, SCFA and gut-repair rate and every coupling into the systemic states; the worst, the SCFA effect on inflammation, had an RSE of 16,865%. One parameter, the Agent A T-cell scale, was classed locally identified in the three-target calibration (RSE 46%) but non-identifiable in the full branch (2,056%), which shows how much the verdict depends on the design assumed.

The interaction itself was invisible to the model’s own outputs. Setting η to −0.5, 0 and +0.5 moved the calprotectin proxy only from 0.650 to 0.644 to 0.641, and the systemic inflammation ratio from 0.9816 to 0.9811 to 0.9808. The stage concluded HOLD and recommended model reduction rather than combination fitting.

### One interaction, six identifiable parameters

The reduced model, AgeQSP v0.9.2-R, keeps only the lead hypothesis. Each on/off rate pair was re-expressed as an amplitude and a time constant (k_on = α/τ, k_off = (1 − α)/τ); the Agent A and Agent B time constants were fixed; the systemic couplings were collapsed into two sensitivity-only coefficients; the Stage-5M observation scales were frozen; H2–H7 were excluded; and η_A×B was locked at 0. Six parameters remain active (Table 2).

**Table 2.** Active parameters of AgeQSP v0.9.2-R under the optimised 20-sample design.

| Parameter | Meaning | Nominal | RSE, 5% CV (%) | RSE, realistic assay noise (%) | Profile-likelihood range |
| --- | --- | --- | --- | --- | --- |
| $\alpha_{\text{ExpA}}$ | Agent A gut-exposure amplitude | 0.667 | 1.5 | 1.8 | Bounded |
| $\alpha_{\text{EngB}}$ | Agent B engraftment amplitude | 0.667 | 1.1 | 0.7 | Bounded |
| $\alpha_{\text{SCFA}}$ | SCFA response amplitude | 0.600 | 7.2 | 3.6 | Bounded |
| $\tau_{\text{SCFA}}$ | SCFA time constant | 12.2 d | 29.6 | 17.1 | 0.53–1.86× nominal |
| $\alpha_{\text{GutRepair}}$ | Gut-repair amplitude | 0.667 | 6.0 | 5.1 | Bounded |
| $\tau_{\text{GutRepair}}$ | Gut-repair time constant | 20.3 d | 28.9 | 28.8 | 0.52–1.78× nominal |

**Table 3.** Enrolment needed to detect a given interaction ratio (CV 0.6, baseline correlation 0.6, two-sided α 0.05, 15% attrition)

| True $R_{int}$ | 80% power: analysable per arm / total enrolled | 90% power: analysable per arm / total enrolled |
| --- | --- | --- |
| 0.65 | 34 / 160 | 45 / 212 |
| 0.70 | 49 / 232 | 66 / 312 |
| 0.75 (base case) | 75 / 356 | 100 / 472 |
| 0.80 | 125 / 592 | 167 / 788 |
| 0.85 | 234 / 1,104 | 314 / 1,480 |
| 0.90 | 557 / 2,624 | 746 / 3,512 |

Realistic noise assumed CVs of 25% for SCFA, 30% for Agent B abundance and the stimulated cytokine response, 15% for T cells and about 45% for calprotectin with duplicate stool samples. All RSEs come from noise-free synthetic data and are design properties, not biological confidence intervals.

The washout carries the information. A greedy D-optimal search over 112 candidate samples chose 20, of which 15 fell in the 28-day washout and none on day 84. With the six parameters, this design had full rank and a condition number of 30.7; a day-84-only design had a condition number of 3,970 and RSEs of 451% for the SCFA amplitude and 10,155% for its time constant. One-at-a-time ±50% perturbation reversed no predicted direction; the day-84 calprotectin proxy ranged from 0.43 to 0.83 around its nominal 0.64.

### A factorial trial that can falsify the lead hypothesis

The model’s design output is a randomised, double-blind, placebo-controlled 2×2 factorial trial, here called A×B, in adults aged about 65–80 with repeated evidence of low-grade inflammation. Participants are allocated 1:1:1:1 to double placebo, Agent A, Agent B or both, at the daily doses used in their monotherapy trials, stratified by site, sex, age band and baseline calprotectin band, for 84 days followed by a 28-day washout. The single primary endpoint is day-84 log faecal calprotectin, analysed as log(calprotectin_84) ∼ A + B + A×B + log(baseline) + strata. The primary estimand is the interaction ratio

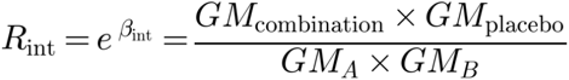

so that R_int < 1 means the combination lowers calprotectin more than multiplicative independence predicts [20]. SCFA, Agent B abundance, stimulated cytokine response, T cells and safety are key secondary endpoints; hsCRP, cytokines, DNA-methylation age, grip strength, 6-minute walk and frailty are exploratory only.

The base case enrols 356. A blinded sample-size re-estimation at about 25% recruitment uses pooled variance only, with an operational cap of 600; 6 of 24 plausible variance scenarios would exceed that cap and trigger redesign. Whether the products make this a clinical trial of an investigational medicinal product in the UK is undetermined and needs regulatory borderline advice.

### A virtual trial found the design’s weak point: the assay floor

A synthetic population of 9,000 people aged 65–80 (median baseline calprotectin 31 mg/kg, median hsCRP 2.4 mg/L, mean adherence 0.92) passed 15 of 15 qualification checks. Simulated trials of 89 randomised per arm, 1,200 replicates per scenario, reproduced the design (Figure 4). In the base case the type I error was 4.5%, power at R_int 0.75 was 80.5%, and the estimate was unbiased (mean 0.750, coverage 95%).

**Figure 4.**
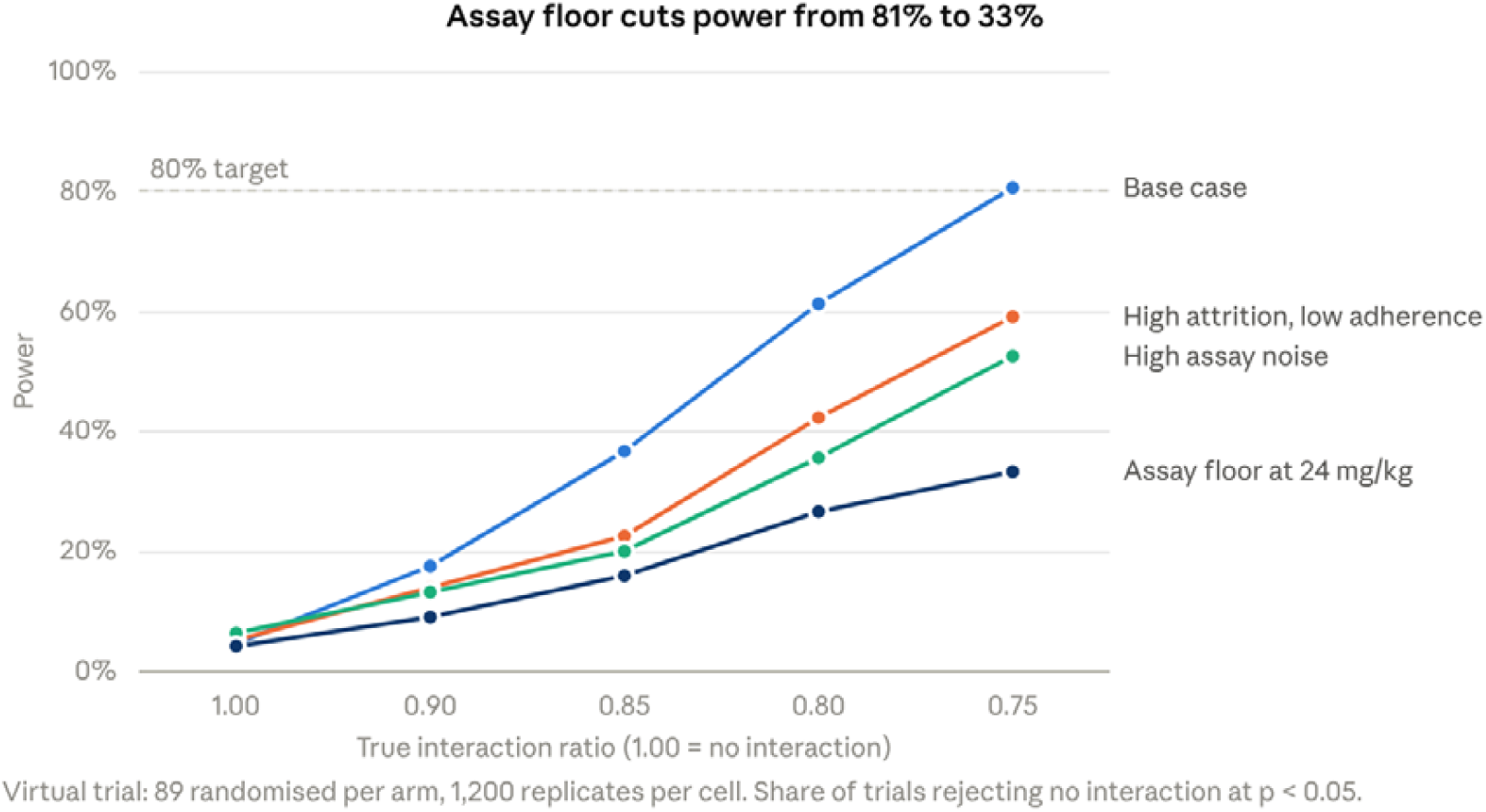
Operating characteristics of the A×B design in a 9,000-person virtual population. Power to reject no interaction (p < 0.05) against the true interaction ratio, for the base case and three stresses; 89 randomised per arm and 1,200 replicates per point. An assay floor at 24 mg/kg, with censored values replaced by the floor, cuts power at a ratio of 0.75 from 81% to 33%. Source: Stage 9A.

Three stresses cost power. Higher assay noise reduced it to 52% and raised the type I error slightly, to 6.3%. High attrition with lower adherence reduced it to 59% and biased the estimate towards no interaction (mean 0.786). The worst case was an assay floor: 37% of simulated baselines lay below 24 mg/kg, and substituting 24 for censored values cut power to 33%, pulled the estimate to 0.895 and dropped coverage to 30%. The stage returned a design condition. A censoring addendum now requires the assay’s limit of quantification to be fixed before lock and censored values to be handled by a prespecified censored-data method, with floor substitution only as a sensitivity analysis, and the locked analysis engine refuses to run if any day-84 value is censored. The addendum has not yet been folded into protocol version 1.0.

### The interaction has no prior

The evidence stage declined to build an informative prior for the interaction. No factorial data exist, and the monotherapy endpoints are too heterogeneous to pool. The primary analysis is frequentist. Three supportive priors on log R_int, normal with mean 0 and SD 0.2, 0.35 or 0.7, give prior probabilities of R_int < 0.75 of 0.075, 0.204 and 0.341. With the planned standard error of 0.102, the chance of a posterior probability of benefit of at least 0.975 is 1.5–2.5% under no interaction and 73–80% at R_int 0.75. All three must be reported, a conclusion that changes across them is labelled prior-sensitive, and monotherapy or synthetic results may never serve as the interaction prior.

### From a locked model to a trial that cannot start without real evidence

After the design, the operator asked the workflow to continue through every remaining stage. It produced sub-stages 9B–9E and 33 further stages (10–42), summarised in Table 4. None used real trial data, because none exists; every stage that needed it stopped at HOLD.

**Table 4.** Stages 9B–42.

| Stages | Purpose | Key result | Verdict |
| --- | --- | --- | --- |
| 9B–9E | Executable SBML and SED-ML for v0.9.2-R | Static checks and replay passed (maximum relative difference $1.8 \times 10^{-16}$); formal libSBML and libSEDML could not run in the environment | HOLD three times |
| 9E-alt, 9E-R | Independent execution route; validation policy 1.1 | Direct MathML execution matched the Python model in 660 state comparisons (maximum relative difference $2.5 \times 10^{-8}$) and 165 output comparisons ($2.7 \times 10^{-8}$); policy changed to accept one qualified route | PASS under the revised policy |
| 10 | Locked primary-analysis engine | On synthetic data a planted interaction of $\beta = -0.220$ was recovered as $-0.197$; unblinding blocked unless database, analysis plan and model hash are locked | PASS |
| 11–12 | Data acquisition, reconciliation, end-to-end dry run | All 7 seeded error classes detected; 24 synthetic participants processed; 6 discrepancies left open | PASS; not ready for sites |
| 13–15 | Activation dossier, evidence ledger, submission bundle | Six activation gates (A1–A6: assay, regulatory route, data capture, eligibility, visit windows, randomisation and unblinding); 16 of 17 submission items open | PASS as tooling; activation HOLD |
| 16–20 | Release simulation, information requests, evidence promotion, import audit | Simulation-labelled evidence closed gates in simulation mode and was rejected in real mode every time | PASS |
| 21–25 | Real-evidence watcher, hash-chained ledger, integrity gate, two-person review, final dry run | 0 of 6 real gates ready; tampered or mislabelled candidates blocked; real-mode dry run blocked | PASS |
| 26–42 | Sponsor pack, monitoring, blinded re-estimation, database lock, interaction estimation, recalibration, final release | 10 framework stages completed; 7 held for evidence or data that do not exist | 10 PASS (framework only), 7 HOLD |

The final state is recorded as a hold at the real-evidence and randomised-data boundary. The interaction parameter η remains 0, H2–H7 remain frozen, the validated archive and analysis engine are pinned by hash, and the last stage records that further PASS verdicts would fabricate evidence that is not available.

Four findings from these stages bear on how much weight the controls can carry.

#### 1. A gate was redefined rather than passed

Formal SBML and SED-ML validation of the reduced model failed to run three times because the validators could not be installed. At the operator’s request, the policy was changed to accept any qualified validation route, and the independent execution route passed. That route’s own report states that it does not replace formal certification, and the released archive is labelled validated under the revised policy. The parent v0.9 model did pass formal libSBML and libSEDML validation in an earlier environment. When we ran libSBML 5.21.2 and libSEDML 2.0.34 on the released archive during manuscript preparation, the SED-ML passed but the SBML had one error: the day-84 treatment-stop event referred to time as an undeclared identifier instead of the SBML time symbol, and the substitute route had simply read it as simulation time. libRoadRunner 2.10.0 refused to load the released file. After a one-element correction it loaded, ran the washout, and matched the Python reference in all 660 state comparisons (maximum relative difference 7.9×10^−8^). The broken event is the one that starts the washout, which carries the model’s identifiability.

#### 2. Simulation detection relied on keywords

In one mixed batch, five fixtures labelled as real-like QA passed as non-simulation evidence, and the batch was rejected only because a sixth file was labelled SIM_ONLY. The token list was extended at Stage 21, but a label is not provenance.

#### 3.Signatures are digests

The signed manifests and reviewer attestations are SHA-256 digests, not cryptographic signatures bound to identities.

#### 4. Framework PASS is not execution

Stages 26–42 were generated as one batch from a shared template, and the ten that passed built runbooks, templates and checks. None executed on trial data.

Throughout, every stage report prohibits clinical efficacy, synergy or rejuvenation claims from simulation outputs, and 15 gate files carry an explicit flag that synthetic or mock data may not update η.

### The same gates in other programmes

Three shorter runs show the same pattern: generation over-reports novelty, and structure can masquerade as mechanism.

#### Ribosome-biogenesis corpus

A literature-only run over one laboratory’s recent publications generated 24 target and combination proposals. A precedent sweep added afterwards found clinical precedent for 7, split verdicts for 2, preclinical precedent for 4, only adjacent precedent for 9 and none for 2. The lead hypothesis, RNA polymerase I inhibition in NPM1-mutated AML, already had clinical support from dactinomycin remissions [21]. The sweep also corrected the run’s own trial design: bulk TP53 variant allele frequency was the wrong stopping rule, because biallelic TP53 loss, visible by single-cell sequencing years before transformation, carries the signal that heterozygous clones do not [22]. Patents and registries were not searched systematically, so the novelty counts are upper bounds; surviving proposals are reported as counts only.

#### NPM1-mutant AML

A sequencing and rescue screen ranked partners for a menin inhibitor and venetoclax-based therapy. Because each partner’s standardised effect was an input and entered the model at a different point, the ranking could reward the entry point rather than the biology. Mechanism-free null archetypes entering at the same points, and a threshold-sensitivity grid, were added before any ranking is reported [23].

#### A second ageing model, IQN-BIOAGE-01

This model, described in a companion preprint [28], couples a stochastic senescent-cell production–removal core [24] to eight latent axes read through 13 blood markers, calibrated on NHANES III and tested on a locked NHANES IV hold-out [26]. Its mortality discrimination matched PhenoAge [25] (Harrell’s C 0.842 for both; ΔC −0.0003, 95% CI −0.0026 to +0.0020). In that model the inflammation axis, one of the reversible axes that track current physiological load, is the largest contributor to variance in the biological-age gap [28]. Together with the AgeQSP validation failure, this is consistent with blood-based clocks weighting reversible physiological change heavily, although both results are model-based and do not establish it. Neither model supports a clock as the primary endpoint of this gut-inflammation combination trial.

#### The gates failed their own tests

Before adopting two amendments to the workflow specification, we ran their acceptance tests on archived runs, and both failed. A model version cited as a case of mechanism cancellation in fact produced its phenomenon in 71 of 300 admissible parameter sets; the original failure had been a scale-referencing error. A proposed scale check, multiplying every rate constant by a common factor, cannot work, because uniform scaling only rescales time and leaves steady states unchanged. Neither amendment was adopted. Separately, when the same architecture was applied to bioprocess interventions, a patent search alone blocked five of six candidates [27].

## Discussion

The workflow’s product was a narrowing, not a recommendation. Seven hypotheses became one interaction; 48 extension parameters became 6; an untestable claim of synergy became a single estimand, R_int, with a design able to reject it. Each step of that narrowing was forced by a gate and recorded, which is what makes it auditable.

The gates changed the science at four points. The evidence gate moved hsCRP from a co-primary to an exploratory endpoint, because neither monotherapy moved systemic CRP. The validation gate kept every clock claim out of the programme after the magnitude failure, so clocks appear in the trial only as exploratory endpoints. The identifiability gate forced the model reduction and showed that the washout, not the end of treatment, identifies the kinetics. And the virtual trial found an assay-floor vulnerability that would have cut power from 81% to 33%; it costs little to fix before a protocol is locked and would be very expensive to find afterwards.

The design is honest about what it can detect. At 356 participants it is powered for an interaction ratio of 0.75, a large interaction. A true ratio of 0.90 would need 2,624. A null result would therefore exclude only a large interaction, not a modest one, and the Agent A main effect on calprotectin, which has no direct trial evidence, was set to neutral in the virtual population. If the trial runs, its most likely informative outcomes are a clear interaction or a clear absence of a large one.

The weakest points were in governance, not modelling. A validation gate that could not be met in the environment was redefined at the operator’s request. The substitute route agreed with the reference model to 10^−8^, yet it accepted a malformed time reference that left the released model unloadable in a standard simulator, which formal validation caught at once. The episode shows both that the person running the workflow can move a gate and why the original gate existed. The audit trail made the change visible; it did not prevent it. Keyword-based detection of simulated evidence let real-looking fixtures through once. And stage counts inflate easily: ten of the last seventeen stages passed while doing nothing on data. A verdict of PASS says a stage met its own criteria, not that the programme advanced.

These observations suggest a minimum for reporting AI-generated combination hypotheses:

1. The corpus, its scope and its declared blind spot.
2. The number of proposals generated before gating, not only the survivors.
3. Direct and precedent evidence for each proposal, with the search scope: literature, registries, patents and pipelines.
4. Parameter provenance and identifiability for the parameters that drive any ranking.
5. The null model of non-interaction, and attribution controls for rankings.
6. The validation tier of each claim, and what evidence would raise it.
7. The claim register, including retractions, and the acceptance-test status of the workflow version used.
8. Every gate that was redefined, waived or overridden, and by whom.
9. The language models used and the points at which humans signed off.

### Limitations

This is one group’s experience, and the AgeQSP record is a single programme run over three days. Outputs depend on the language models used, which change between versions. Nearly all combination results are Tier 0: kinetic parameters are assumed, the relative standard errors rest on assumed observation noise and noise-free synthetic data, and the virtual population’s variability is assumed. Agent B’s calprotectin anchor is one small, short study in a population older than the planned trial, which enrols from 65. The scripts behind the sample-size table, the realistic-noise identifiability check, the virtual trial and the Bayesian operating characteristics are not in the archive, so those results cannot yet be regenerated from it. Minor inconsistencies remain between stage files, for example in the timing of the post-treatment IL-6 sample (days 85 and 87 in the optimised design, day 86 in the calendar). Finally, no hypothesis in this paper has been tested in people.

### Conclusion

An agentic workflow can now generate combination hypotheses, models and trial designs faster than a team can check them. In this programme the useful output was the record of what the gates removed and why: one falsifiable interaction, a design that can test it, a known weak point in its assay, and a hold at the point where only real data can advance it.

## Methods

### Workflow

All AgeQSP work ran under version 2.21 of IQANOVA’s AI-QSP workflow specification, a staged specification executed by LLM agents with file, code and web. Each stage returned its inputs, work, provenance, assumptions, blocking conditions, deliverables and exactly one verdict (PASS, HOLD or BLOCKED). The operator issued short control commands and one governance change (Results). The specification’s prompt text is proprietary and not reproduced; its stage map and gates are described in Box 1.

### Model

AgeQSP is a system of ordinary differential equations on dimensionless latent states (0–1) with explicit observation models for each biomarker. Drug pharmacokinetics use a two-compartment subcutaneous model for semaglutide and a one-compartment oral model for metformin, with scheduled dosing events. The Agent A–Agent B extension adds first-order exposure and engraftment states coupled multiplicatively to parent inflammation and immune-decline states. Models were encoded in SBML Level 3 Version 2 with SED-ML Level 1 Version 4 simulation experiments and COMBINE/OMEX archives [6,7]; Python implementations served as reference.

### Agent coding

To protect pending intellectual property, the combination partners are identified by codes in this version: Agent A, a food-derived bioactive ingredient; Agent B, a single-strain probiotic; Agent C, a widely used nutritional supplement; Agent D, a supplement with proposed mitochondrial effects; Agent E, an immunomodulatory drug class (drug and dose not fixed); Agent F, a synbiotic; and Agent G, a micronutrient. Their doses, the calibration target values and the trials used to calibrate them are withheld, and citations to those trials are marked “reference withheld”. Fitted and assumed parameters for Agents A and B are reported under coded names (Table 2). The semaglutide and metformin branches are named because their calibration and validation use published single-agent data. The workflow, model structure, identifiability analysis, trial design and virtual-trial results can be followed without the identities, although the doses and calibration targets come from agent-specific trials. The identities will be disclosed in a later version.

### Calibration and validation

Parameters were fitted by least squares to aggregate published targets, some digitised from figures, and every parameter carries a provenance label. Validation used models frozen before comparison, with no refitting; metrics were sign concordance, coverage by the reported 95% confidence interval and RMSE.

### Sensitivity and identifiability

Global sensitivity used Latin hypercube sampling (N = 3,000; ±30% around nominal) with Spearman rank correlation. Local identifiability used Fisher information from central finite differences (±5% in log coordinates), reporting rank, condition number, relative standard errors and pairwise correlations under assumed observation noise (5% normalised error unless stated). Profile likelihoods were re-optimised on synthetic pseudo-data with a threshold of ΔRSS = 3.84, and robustness was tested one parameter at a time at ±50%. The sampling design was chosen by greedy constrained D-optimality over 112 candidates (14 days × 8 endpoints), with at most 4 samples per endpoint and per day and at least 6 in the washout.

### Evidence synthesis

STEP 1–3 week-68 effects were pooled with a normal–normal random-effects model, half-normal priors on between-trial SD (5 percentage points for weight; 0.25 on the log scale for CRP) and numerical integration over the SD. For the trial, three supportive priors on log R_int (normal, mean 0, SD 0.2, 0.35 or 0.7) were specified before any data, and their operating characteristics were computed at the planned standard error.

#### Sample size

With total coefficient of variation CV, baseline–outcome correlation ρ and n analysable participants per arm, the standard error of the log interaction is

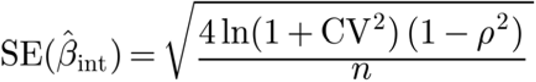

which gives 0.102 at CV 0.6, ρ 0.6 and n = 75, and 80% power at R_int 0.75 with two-sided α 0.05. Enrolment allows for 15% attrition.

#### Virtual population and virtual trial

Nine thousand synthetic participants were drawn with age from a truncated normal (mean 72.5, SD 4.1, 65–80), baseline calprotectin and hsCRP from log-normals, adherence from a beta distribution, and the six active parameters from correlated log- or logit-normal distributions centred on nominal values. Each virtual trial randomised 89 per arm and analysed day-84 log calprotectin by ANCOVA with the interaction term, baseline, age and sex (the analysis plan adjusts for the randomisation strata instead), with 1,200 replicates per scenario.

#### Executable validation

The v0.9 extension was validated with libSBML 5.21.2 and libSEDML 2.0.34 and a parent zero-switch replay (tolerance 10^−6^). For v0.9.2-R, where the validators could not be installed, an independent route parsed the SBML MathML directly, integrated all 49 rate-rule states with SciPy to day 112, and compared states and SED-ML outputs with the Python reference (tolerance 10^−6^). During manuscript preparation we ran libSBML 5.21.2, libSEDML 2.0.34 and libRoadRunner 2.10.0 on the released archive, corrected the single error found in a copy by replacing the time identifier with the SBML time symbol, and compared the copy with the Python reference at the 660 state–time points used by the independent route (integrator tolerances 10^−12^ absolute and 10^−10^ relative). Release archives and every later stage are pinned by SHA-256, and each stage manifest records the hash of the previous one.

## Data and code availability

Coded versions of the following will be deposited at IQANOA ATLAS (www.iqanova.org) the AgeQSP v0.9.2-R archive as released, the Python reference models; the locked primary-analysis engine; the virtual population; and the stage manifests. In the coded files every model identifier and name that would identify an agent is replaced by the codes used here, and annotations, notes and ontology terms that would do the same are removed; the coded archives therefore have their own hashes. The uncoded archives, the v0.9 all-hypotheses model, the hypothesis register, the evidence ledger and the contradiction–support map will be deposited when the agent identities are disclosed. All trial inputs are published aggregate data; no individual-level human data were used. The workflow specification is proprietary to IQANOVA Ltd.

## Declarations

### Ethics

This study used only published aggregate data and synthetic data. No participants were recruited and no ethics approval was required. The A×B design has not been submitted for regulatory or ethics review.

### Use of generative AI

LLM agents executed the workflow stages, wrote and ran code, and drafted stage reports, as described in Methods IQANIOVA AI-QSP platform. LLMs also assisted drafting of this manuscript. The authors checked the reported numbers against the source files and take full responsibility for the content. No AI system is an author.

### Funding

Innovate UK AI Champions Frontier AI programme (ATLAS-AI), IQANOVA LTD

### Competing interests

I.G. is a director of IQANOVA Ltd, which develops the workflow commercially. IQANOVA Ltd holds UKIPO patent applications relating to AI-assisted model-informed drug development.

### Author contributions

IG run the pipeline, BD edited the manuscript IVG provide admin support and funding. AI assistance is described above and is not an author contribution.

## Supplementary information

**Table S1.** Full hypothesis register, model encoding and therapeutic scenarios, coded (predictions and falsifiers are in Table 1)

| ID | Class | Mechanism as generated | Key endpoints | Sources | Encoded model path (v0.9) | Scenario and stage decision |
| --- | --- | --- | --- | --- | --- | --- |
| H1 | Nutritional/biologic adjunct combination | Agent A: immune-cell and inflammatory modulation; Agent B: gut inflammatory reduction. Proposed interaction through microbiome/SCFA and gut-to-systemic inflammatory coupling | Faecal calprotectin; hsCRP; IL-6; TNF-α; stimulated cytokine responses; T-cell subsets; shotgun microbiome; acetate/butyrate; GI tolerability | [18, 19] and agent-specific studies (references withheld) | A_gut, B_engraft, SCFA, GutRepair → inflammation, immune decline | TS1: ages 65–80, stable, repeat hsCRP about 2–10 mg/L; 12 weeks; ADVANCE TO TRIAL DESIGN |
| H2 | Multi-axis geroscience combination | Gut/immune repair plus systemic inflammation and DNAm-ageing effects; tests whether gut-inflammatory improvement adds to Agent C's own reported effects | H1 endpoints + DNA-methylation clocks, frailty index, infections, falls | [18] and agent-specific studies (references withheld) | H1 + Agent C signal → inflammation, mitochondria, nutrient sensing | TS2: after TS1 proof of mechanism; 6–12 months; CONDITIONAL |
| H3 | Gut-immune + mitochondrial/muscle combination | Agents A and B target gut inflammation and immune tone; Agent D is proposed to improve mitochondrial function and muscle endurance | Muscle endurance; physical function; mitochondrial and metabolic markers; CRP; calprotectin; frailty | Agent-specific studies (references withheld) | H1 + Agent D signal → mitochondria | TS3: reduced physical function; several months; CONDITIONAL |
| H4 | Drug + gut-immune adjuncts | Incretin metabolic/weight pathway plus reduction of residual gut/systemic inflammaging; not enhanced weight loss | Weight; HOMA-IR; CRP; calprotectin; immune subsets; GI adverse events; organ-age measures exploratory | Agent-specific studies (references withheld) | Inherited drug branch + H1 | TS4: obesity or metabolic dysfunction; 6–12 months; EXPLORATORY |
| H5 | Drug + gut-metabolic adjuncts | Biguanide gut/systemic metabolic signalling plus Agent A/B modulation of microbiota, barrier inflammation and SCFA production | HOMA-IR; GDF15; weight; calprotectin; SCFA; microbiome; GI adverse events | [18] and agent-specific studies (references withheld) | Inherited drug branch + H1 | No scenario; 96-week simulation only |
| H6 | Pharmacologic immune-ageing + gut-immune adjuncts | Agent E may enhance antiviral immunity; Agent A alters cytokine/T-cell responses; Agent B reduces gut inflammation | Immune-response markers; infection outcomes; T-cell subsets; CRP/calprotectin; safety labs | Agent-specific studies (references withheld) | H1 + Agent E signal → immune decline and two other systemic states | TS5: infection risk; seasonal course; LOWER PRIORITY |
| H7 | Microbiome-conditioning combination | Agent A and the prebiotic component of Agent F can shift target taxa; Agent G may support host physiology | Target-taxon absolute abundance; SCFA; barrier and inflammatory markers; micronutrient status | [18] and agent-specific studies (references withheld) | A_gut + F_engraft + prebiotic substrate + Agent G signal → SCFA, gut repair, inflammation, immune decline | No scenario; 12-week simulation only |

**Table S2.** Contradiction–support map (AgeQSP v0.9), coded.

| Evidence item | Supports | Contradiction or limitation | Resolution in v0.9 | Gate |
| --- | --- | --- | --- | --- |
| Agent A older-adult RCT (reference withheld) | Agent A immune/inflammation branch | Short trial; no ageing endpoint; plasma CRP/IL-6 not clearly superior to placebo | Encode Agent A couplings as ASSUMED until calibrated | PASS HYPOTHESIS |
| Agent B older-adult RCT (reference withheld) | Gut-inflammation/probiotic branch | CRP trend non-significant; calprotectin improved | Encode gut/SCFA branch; do not force a systemic CRP effect | PASS HYPOTHESIS |
| 2025 probiotic/prebiotic/synbiotic meta-analysis [18] | Microbiome/SCFA/inflammatory priors | High intervention heterogeneity | Use as prior range, not calibration target | PASS WITH CAUTION |
| Related multi-component supplement study (reference withheld) | Microbiome-conditioning H7 branch | Taxa shifted; metabolite, barrier and inflammatory markers not improved | Keep H7 as a microbiome-conditioning hypothesis; no barrier-benefit claim | PASS WITH CONTRADICTION |
| Clock evidence for the add-on agents (references withheld) | H2 and H7 add-on priors | Small or post hoc effects | Encode H2/H7 add-ons only; clocks exploratory | PASS HYPOTHESIS |
| Agent D RCT (reference withheld) | Mitochondrial H3 branch | Primary functional endpoints non-significant | Encode Agent D as hypothesis; retain the negative primary result | PASS HYPOTHESIS |
| Agent E evidence (reference withheld) | H6 antiviral-immune branch | Clinical infection benefit not confirmed in a large trial | Sensitivity-only immune branch; lower priority | PASS WITH WARNING |
| Direct AxB factorial data | Would identify $\eta_{A \times B}$ | Not found | Interaction stays ASSUMED_SENSITIVITY_ONLY | BLOCK CALIBRATION |

**Table S3.**
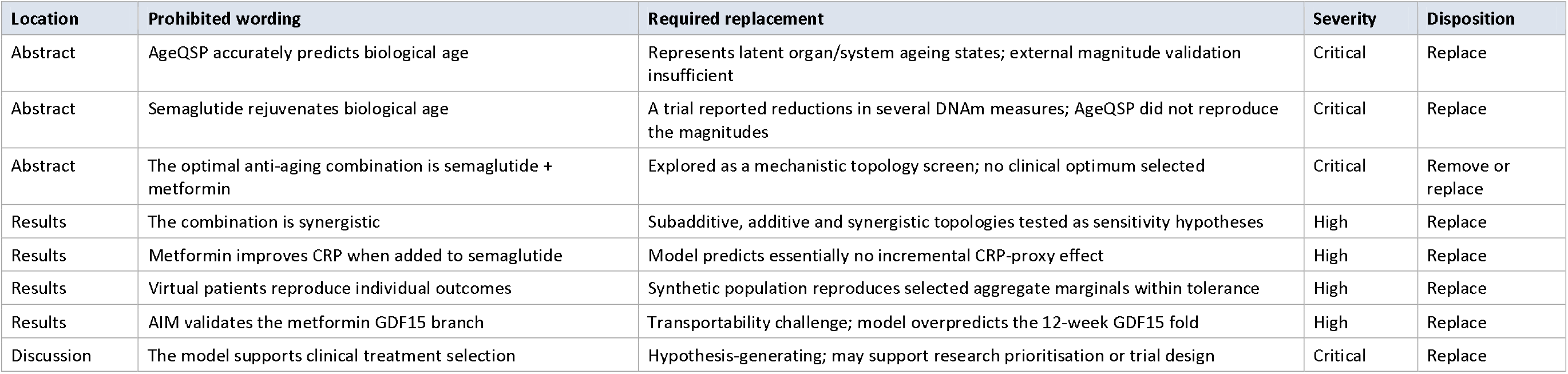
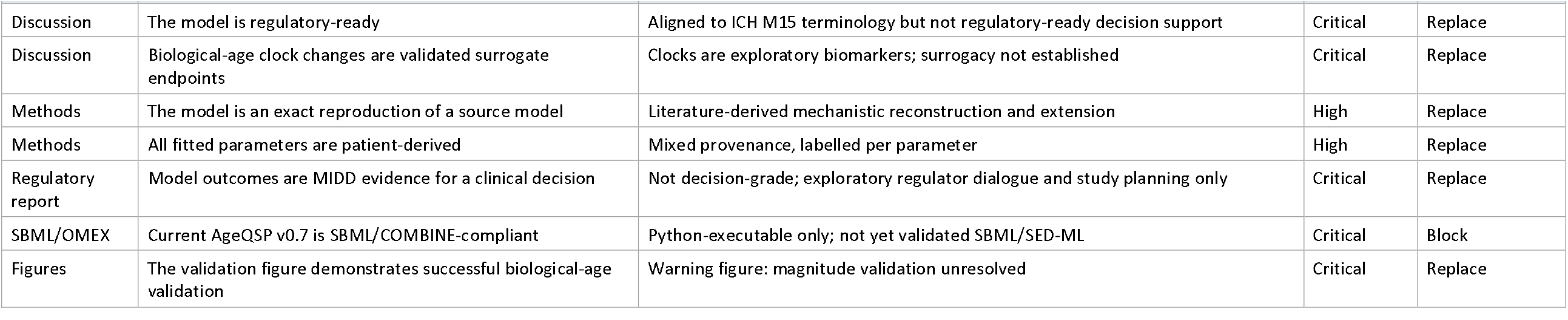
Claim-warning table applied to the AgeQSP v0.8 report (Stage 12A)

## References

1. Sorger PK, Allerheiligen SRB, Abernethy DR, Altman RB, Brouwer KLR, Califano A, et al. Quantitative and systems pharmacology in the post-genomic era: new approaches to discovering drugs and understanding therapeutic mechanisms. An NIH White Paper by the QSP Workshop Group. Bethesda (MD): National Institutes of Health; 2011. Available from: https://www.nigms.nih.gov/training/documents/systemspharmawpsorger2011.pdf

2. Chelliah V, Lazarou G, Bhatnagar S, Gibbs JP, Nijsen M, Ray A, et al. Quantitative systems pharmacology approaches for immuno-oncology: adding virtual patients to the development paradigm. Clin Pharmacol Ther. 2021;109(3):605–618. doi:10.1002/cpt.1987.

3. Androulakis IP, Cucurull-Sanchez L, Kondic A, Mehta K, Pichardo C, Pryor M, et al. The dawn of a new era: can machine learning and large language models reshape QSP modeling? J Pharmacokinet Pharmacodyn. 2025;52(4):36. doi:10.1007/s10928-025-09984-5.

4. Zhavoronkov A, Naumov V, Sidorenko D, Aliper A, Aladinskiy V, Hasani R, et al. An open benchmark and language models for AI in aging biology. Cell. 2026;189(19):5980–5994.e8. doi:10.1016/j.cell.2026.08.026.

5. Bischoff-Ferrari HA, Gängler S, Wieczorek M, Belsky DW, Ryan J, Kressig RW, et al. Individual and additive effects of vitamin D, omega-3 and exercise on DNA methylation clocks of biological aging in older adults from the DO-HEALTH trial. Nat Aging. 2025;5(3):376–385. doi:10.1038/s43587-024-00793-y.

6. Goryanin I, Damms B, Goryanin I. A systems pharmacology model of aging identifies optimal combination therapies with secondary benefits on weight loss and metabolic health. CPT Pharmacometrics Syst Pharmacol. 2026;15(8). doi:10.1002/psp4.70322.

7. Keating SM, Waltemath D, König M, Zhang F, Dräger A, Chaouiya C, et al. SBML Level 3: an extensible format for the exchange and reuse of biological models. Mol Syst Biol. 2020;16(8). doi:10.15252/msb.20199110.

8. Bergmann FT, Adams R, Moodie S, Cooper J, Glont M, Golebiewski M, et al. COMBINE archive and OMEX format: one file to share all information to reproduce a modeling project. BMC Bioinformatics. 2014;15:369. doi:10.1186/s12859-014-0369-z.

9. American Society of Mechanical Engineers. V&V 40-2018: Assessing credibility of computational modeling through verification and validation: application to medical devices. New York: ASME; 2018.

10. Kuemmel C, Yang Y, Zhang X, Florian J, Zhu H, Tegenge M, et al. Consideration of a credibility assessment framework in model-informed drug development: potential application to physiologically-based pharmacokinetic modeling and simulation. CPT Pharmacometrics Syst Pharmacol. 2020;9(1):21–28. doi:10.1002/psp4.12479.

11. Li Y, Xu X, Sehgal R, He X, Zheng Y, Chen X, Xu K. Effect of caloric restriction on organ-specific biological aging in a randomized clinical trial. Clin Nutr. 2026;60:106625. doi:10.1016/j.clnu.2026.106625.

12. Wilding JPH, Batterham RL, Calanna S, Davies M, Van Gaal LF, Lingvay I, et al. Once-weekly semaglutide in adults with overweight or obesity. N Engl J Med. 2021;384(11):989–1002. doi:10.1056/NEJMoa2032183.

13. Rubino D, Abrahamsson N, Davies M, Hesse D, Greenway FL, Jensen C, et al. Effect of continued weekly subcutaneous semaglutide vs placebo on weight loss maintenance in adults with overweight or obesity: the STEP 4 randomized clinical trial. JAMA. 2021;325(14):1414–1425. doi:10.1001/jama.2021.3224.

14. Garvey WT, Batterham RL, Bhatta M, Buscemi S, Christensen LN, Frias JP, et al. Two-year effects of semaglutide in adults with overweight or obesity: the STEP 5 trial. Nat Med. 2022;28(10):2083–2091. doi:10.1038/s41591-022-02026-4.

15. Verma S, Bhatta M, Davies M, Deanfield J, Garvey WT, Jensen C, et al. Effects of once-weekly semaglutide 2.4 mg on C-reactive protein in adults with overweight or obesity (STEP 1, 2, and 3): exploratory analyses of three randomised, double-blind, placebo-controlled, phase 3 trials. EClinicalMedicine. 2023;55:101737. doi:10.1016/j.eclinm.2022.101737.

16. Perreault L, Davies M, Frías JP, Laursen PN, Lingvay I, Machineni S, et al. Changes in glucose metabolism and glycemic status with once-weekly subcutaneous semaglutide 2.4 mg among participants with prediabetes in the STEP program. Diabetes Care. 2022;45(10):2396–2405. doi:10.2337/dc21-1785.

17. Corley MJ, Dwaraka VB, Pang APS, Labbato D, Smith R, Eckard AR, et al. Semaglutide slows epigenetic aging in a randomized trial of HIV-associated lipohypertrophy. Nat Commun. 2026;17:6606. doi:10.1038/s41467-026-72861-3.

18. Kolnes KJ, Møller PM, Kruse R, Christensen MMH, Kjøbsted R, Hey-Mogensen M, et al. Metformin increases glycolysis and the stress-induced cytokine GDF15 but not FGF21 in humans. Front Endocrinol (Lausanne). 2026;17:1797525. doi:10.3389/fendo.2026.1797525.

19. Gao F, Li C, Wang Y, et al. Growth differentiation factor 15 is not associated with glycemic control in patients with type 2 diabetes mellitus treated with metformin: a post-hoc analysis of the AIM study. BMC Endocr Disord. 2022;22(1):256. doi:10.1186/s12902-022-01176-3.

20. Berthon BS, Williams EJ, Williams LM, Budden KF, Hiles SA, Bartlett NW, Wood LG. Oral lactoferrin reduces systemic inflammation, enhances anti-viral responses and modulates immune cell profiles: a randomised controlled trial in healthy, older adults. Br J Nutr. 2026;135(9):929–943. doi:10.1017/S000711452610631X.

21. Lazou-Ahrén I, Björklund M, Molin G, Xu J, Önning G, Elmståhl S, Jeppsson B. Probiotic-reduced inflammaging in older adults: a randomized, double-blind, placebo-controlled trial. Probiotics Antimicrob Proteins. 2025;17(5):3429–3439. doi:10.1007/s12602-024-10310-7.

22. Zhuang K, Luo H, Zeng M, Chan SCL, Gong M, Wang Y. Effects of probiotics, prebiotics, and synbiotics on gut microbiota in older adults: a systematic review and meta-analysis of randomized controlled trials. Nutr J. 2025;24:147. doi:10.1186/s12937-025-01218-1.

23. Konstanti P, van Splunter M, van den Brink E, Belzer C, Nauta A, van Neerven RJJ, Smidt H. The effect of nutritional intervention with lactoferrin, galactooligosaccharides and vitamin D on the gut microbiota composition of healthy elderly women. Nutrients. 2022;14(12):2468. doi:10.3390/nu14122468.

24. Suzuki N, Murakoshi M, Ono T, Morishita S, Koide M, Bae MJ, et al. Effects of enteric-coated lactoferrin tablets containing Lactobacillus brevis subsp. coagulans on fecal properties, defecation frequency and intestinal microbiota of Japanese women with a tendency for constipation: a randomized placebo-controlled crossover study. Biosci Microbiota Food Health. 2013;32(1):13–21. doi:10.12938/bmfh.32.13.

25. Neyrinck AM, Rodriguez J, Taminiau B, Amadieu C, Herpin F, Allaert FA, et al. Improvement of gastrointestinal discomfort and inflammatory status by a synbiotic in middle-aged adults: a double-blind randomized placebo-controlled trial. Sci Rep. 2021;11(1):2627. doi:10.1038/s41598-020-80947-1.

26. Liu S, D’Amico D, Shankland E, Bhayana S, Garcia JM, Aebischer P, et al. Effect of urolithin A supplementation on muscle endurance and mitochondrial health in older adults: a randomized clinical trial. JAMA Netw Open. 2022;5(1). doi:10.1001/jamanetworkopen.2021.44279.

27. Mannick JB, Teo G, Bernardo P, Quinn D, Russell K, Klickstein L, Marshall W, Shergill S. Targeting the biology of ageing with mTOR inhibitors to improve immune function in older adults: phase 2b and phase 3 randomised trials. Lancet Healthy Longev. 2021;2(5)–e262. doi:10.1016/S2666-7568(21)00062-3.

28. ClinicalTrials.gov. Effects of a synbiotic supplement on metabolic & cognitive outcomes among older adults. ClinicalTrials.gov identifier: NCT06845995 [Internet]. Bethesda (MD): National Library of Medicine; [cited 2026 Sep 29]. Available from: https://clinicaltrials.gov/study/NCT06845995.

29. National Institutes of Health, Office of Dietary Supplements. Probiotics: fact sheet for health professionals [Internet]. Bethesda (MD): National Institutes of Health; updated 2025 Mar 25 [cited 2026 Sep 29]. Available from: https://ods.od.nih.gov/factsheets/Probiotics-HealthProfessional/

30. Bliss CI. The toxicity of poisons applied jointly. Ann Appl Biol. 1939;26(3):585–615. doi:10.1111/j.1744-7348.1939.tb06990.x.

31. Falini B, Brunetti L, Martelli MP. Dactinomycin in NPM1-mutated acute myeloid leukemia. N Engl J Med. 2015;373(12):1180–1182. doi:10.1056/NEJMc1509584.

32. Kennedy AL, Myers KC, Bowman J, Gibson CJ, Camarda ND, Furutani E, et al. Distinct genetic pathways define pre-malignant versus compensatory clonal hematopoiesis in Shwachman-Diamond syndrome. Nat Commun. 2021;12:1334. doi:10.1038/s41467-021-21588-4.

33. Goryanin I, et al. A stress-testing quantitative systems pharmacology workflow for prioritizing experiments in NPM1-mutant acute myeloid leukemia. Manuscript submitted for publication; 2026.

34. Karin O, Agrawal A, Porat Z, Krizhanovsky V, Alon U. Senescent cell turnover slows with age providing an explanation for the Gompertz law. Nat Commun. 2019;10:5495. doi:10.1038/s41467-019-13192-4.

35. Levine ME, Lu AT, Quach A, Chen BH, Assimes TL, Bandinelli S, et al. An epigenetic biomarker of aging for lifespan and healthspan. Aging (Albany NY). 2018;10(4):573–591. doi:10.18632/aging.101414.

36. Kwon D, Belsky DW. A toolkit for quantification of biological age from blood chemistry and organ function test data: BioAge. Geroscience. 2021;43(6):2795–2808. doi:10.1007/s11357-021-00480-5.

37. Goryanin I, et al. Gated AI-assisted modelling of bioprocess interventions. Manuscript in preparation; 2026.

